# The auditory P2 evoked by speech sounds consists of two separate subcomponents

**DOI:** 10.1101/2023.06.30.547226

**Authors:** Kurt Steinmetzger, André Rupp

## Abstract

The P2 component of the auditory evoked potential is commonly thought to reflect acoustic stimulus properties as well as prior exposure to the materials, rather than change-related cortical activity. Here, we challenge this view by showing that the P2 is strongly increased in response to voice pitch changes with a stepwise pattern compared to changes in dynamic pitch contours typical for natural speech, and also reflects the magnitude of these pitch changes. Furthermore, it is demonstrated that neither the P2 nor any other component are affected by the harmonicity of the materials. Despite no prior exposure, artificially created inharmonic versions of the speech materials elicited similar activity throughout auditory cortex. This suggests that so-called harmonic template neurons observed in animal studies are either absent or do not exist in sufficient number in human auditory cortex to detect their activity extracranially. Crucially, both morphology and source reconstructions of the EEG data showed that the P2 appears to consist of two separate subcomponents. Whereas source activity for the “P2a” was strongest in right auditory cortex, the subsequent “P2b” included generators spread across auditory cortex and association areas, bilaterally. The two subcomponents thus likely reflect processing at different stages of the auditory pathway.

## 1. Introduction

Although it is part of the P1-N1-P2 complex of transient responses and therefore ubiquitous in recordings of auditory evoked cortical activity, functional significance, cortical generators, and morphology of the auditory P2 component have remained elusive. Other components have been studied much more thoroughly, especially the preceding N1, partly because its large amplitude and short duration facilitate the localisation of its sources (e.g., Krumbholz et al., 2003; Näätänen et al., 1987; Roberts et al., 1996). Traditionally, the P2 is thought to reflect the acoustic characteristics of the stimulus materials, whereas the N1 is taken to represent change-related activity (Näätänen et al., 1987). While an N1 is elicited by both sound onsets and offsets, the P2 is only observed following sound onset (Hari et al., 1987). The P2 has been shown to increase with spectral complexity, with the largest amplitudes observed for sounds containing multiple adjacent harmonics, as is typical for musical and speech sounds (Shahin et al., 2005). Moreover, P2 amplitudes in response to harmonic sounds have been found to increase further with repeated stimulus exposure, suggesting that the P2 also reflects familiarity with specific types of sounds (Sheehan et al., 2005; Tremblay et al., 2014). However, recent findings showed that the P2 is also enhanced in response to pitch changes in speech and music (Andermann et al., 2021; Steinmetzger et al., 2022a; Steinmetzger et al., 2022b), implying that it reflects the processing of acoustic changes too.

Regarding the cortical generators of the P2, the findings have also been inconsistent. MEG studies using a dipole-based approach consistently localised the P2 to the lateral part of Heschl’s gyrus, slightly anterior and medial to the N1, irrespective of whether speech or non-speech stimuli were used (Hari et al., 1987; Pantev et al., 1996; Ross et al., 2009; Tiitinen et al., 1999). However, intracerebral recordings (Godey et al., 2001) and recent fMRI-based dipole source localisations of MEG data (Benner et al., 2023) suggested separate sources in planum temporale as well as slightly anterior to Heschl’s gyrus. For EEG data obtained from unilateral cochlear implant (CI) users with preserved contralateral normal hearing, in contrast, a single dipole source of the P2 was localised to the planum temporale (Steinmetzger et al., 2022b). Lastly, distributed MEG source reconstructions of the P2 revealed broadly distributed, right-lateralised activity in auditory areas in response to speech (Coffey et al., 2017).

In terms of the P2 morphology an interesting feature is that it frequently contains two separate peaks. Although most studies did not explicitly discuss this characteristic (Andermann et al., 2017; Bertoli et al., 2011; Steinmetzger et al., 2020; Steinmetzger et al., 2022b; Tiitinen et al., 1999; Tremblay et al., 2014), others have referred to it as ‘distinct second peak’ (Ross et al., 2009) or ‘splitting P2’ (Davis et al., 1966). While this feature does not appear to reflect the type of stimulus material used, there is some evidence that it is more pronounced in middle-aged and older subjects (Bertoli et al., 2011; Ross et al., 2009). However, it has not been investigated yet whether the two peaks might represent separate P2 subcomponents generated in different cortical areas and reflecting different functional processes, demonstrating how little is known about this component.

Prompted by the large, double-peaked P2s observed in response to voice pitch changes in our previous work (Steinmetzger et al., 2020; Steinmetzger et al., 2022a), we here studied the P2 in more detail. Firstly, we compared the effects of pitch change magnitude and the type of pitch change. It has recently been shown that the P2 amplitude reflects the magnitude of pitch changes in musical sequences (Andermann et al., 2021), but it remains unclear if the P2 is also affected by the context in which these changes occur and how both effects compare. Specifically, participants were presented with sequences of speech sounds consisting of stimuli that either had a static pitch or dynamically varying pitch contours typical for natural speech, resulting in stepwise pitch changes confined to the transitions between stimuli or continuous pitch changes, respectively. Larger P2 amplitudes at stimulus onset were expected for stepwise pitch changes due to their greater saliency, despite a similar pitch change magnitude for both types of pitch changes.

Secondly, it was evaluated whether the P2 amplitude is affected by the harmonicity of the stimulus materials, i.e., the property that the frequencies of the spectral components are integer multiples of the fundamental frequency (*F*0). As the P2 is enhanced for sounds with multiple harmonically-related spectral components and also appears to reflect the familiarity with the materials, one would expect larger amplitudes for natural harmonic speech sounds as compared to artificially-created inharmonic versions of these. Indeed, so-called harmonic template neurons, which preferentially fire in response to harmonic sounds, have been observed across the auditory cortex of marmoset monkeys (Feng et al., 2017; Wang, 2018) and to a lesser extent also in the rabbit midbrain (Su et al., 2020). Yet, it is unclear whether such neurons also exist in sufficient number in the human auditory cortex to detect enhanced responses to harmonic sounds extracranially using EEG. Furthermore, it is unknown if the presence of these neurons is confined to the auditory cortex or whether harmonic sounds also elicit larger responses in auditory association cortex.

Finally, we sought to determine if the P2 consists of two separate subcomponents, as suggested by the double-peaked morphology. It was therefore tested if the cortical generators of the two peaks in the waveforms differ and whether they might be evoked at different stages of the auditory processing hierarchy. Specifically, while the first peak was expected to be generated mainly in the auditory cortex, the second peak was assumed to have a broader distribution and to include generators in auditory association areas in the superior temporal cortex too. In contrast to the majority of studies concerned with localising the sources of the P2, distributed source reconstructions were used to be able to estimate the spatial extent of activity.

## 2. Material and methods

### 2.1. Participants

Twenty subjects (9 females, 11 males; mean age 23 years, standard deviation 2.8 years) were tested and paid for their participation. They were all right-handed and reported no history of neurological or psychiatric illnesses. All participants used German as their main language and had audiometric thresholds of less than 20 dB hearing level (HL) at octave frequencies between 125 and 8000 Hz. All subjects gave written consent prior to the experiment and the study was approved by the local research ethics committee (Medical Faculty, University of Heidelberg).

### 2.2. Stimuli

The stimulus materials were the same as in Steinmetzger et al. (2020), where the data were pooled across conditions for analysis. The experiment comprised five different stimulus conditions, four with discrete spectral components and speech-shaped noise. The stimuli with discrete spectral components were based on recordings from the EUROM database (Chan et al., 1995), consisting of five-to six-sentence passages read by 16 different male talkers. Using methods as previously described (Green et al., 2013; Steinmetzger et al., 2015), the *F*0 contours of the sixteen passages were extracted and interpolated through unvoiced and silent periods to generate continuous *F*0 contours.

For the first stimulus condition (*Static F0 – Harmonic*), the log-transformed distribution of the *F*0 values for each individual talker was divided into 12 quantiles and used to generate a set of 192 1-s harmonic complex tones with static pitch contours (16 talkers x 12 quantiles). The complexes were synthesised with equal-amplitude components in sine phase and normalised to a median *F*0 of 100 Hz. To produce the second condition (*Dynamic F0 – Harmonic*), the 16 original pitch contours were used to generate harmonic complexes with dynamically varying pitch tracks. The first 12 s of each tone complex were selected and divided into consecutive 1-s segments. For these two conditions, the frequencies of all component tones were integer multiples of the *F*0 and thus harmonically related. Additionally, inharmonic equivalents of the first two conditions were produced by shifting the frequencies of all component tones by 25% of the median *F*0 (*Static F0 – Inharmonic & Dynamic F0* – *Inharmonic*). This procedure renders the stimuli inharmonic and reduces their pitch strength (Roberts et al., 2001), but leaves all other acoustic properties largely unchanged (Steinmetzger et al., 2023). The components were shifted by 25% as this value was shown to maximise the degree of inharmonicity for tone complexes with a fixed pitch (Roberts et al., 2010). For half the stimuli the shift was applied upwards, for the other half it was applied downwards. A fifth condition (*Speech-shaped noise*), in which the stimuli contained no discrete spectral components and hence no pitch, was based on 192 different 1-s segments of white noise.

All stimuli had a sampling rate of 48 kHz and their spectra were shaped to have a similar long-term average speech spectrum, as described in Steinmetzger et al. (2020). After applying 25-ms Hann-windowed on- and offset ramps, all stimuli were adjusted to have the same root-mean-square level. Example stimuli of all five conditions are shown in Fig. 1A. For the waveforms depicted in the upper row, it is apparent that only the harmonic stimuli have periodic waveforms, while they are less regular for the inharmonic conditions, and completely aperiodic for speech-shaped noise. The narrow-band spectrograms in the middle row demonstrate that the spectra of the stimuli are indeed very similar, despite the markedly different waveforms. To visualise the different degrees of stimulus periodicity, spectrographic representations of summary autocorrelation functions (SACFs; Meddis and Hewitt, 1991; Meddis and O’Mard, 1997; for computational details see Steinmetzger et al., 2020) are shown in the bottom row. While the first peak in these SACF spectrograms represents the *F*0 contours of the stimuli, the height of this peak may be interpreted as a measure of pitch strength (Yost et al., 1996). In line with this notion, the peak around 10 ms is noticeably more pronounced for the harmonic stimuli compared to the inharmonic equivalents. Due to the lack of any temporal regularity, there was no such peak at all for speech-shaped noise.

**Figure 1.**
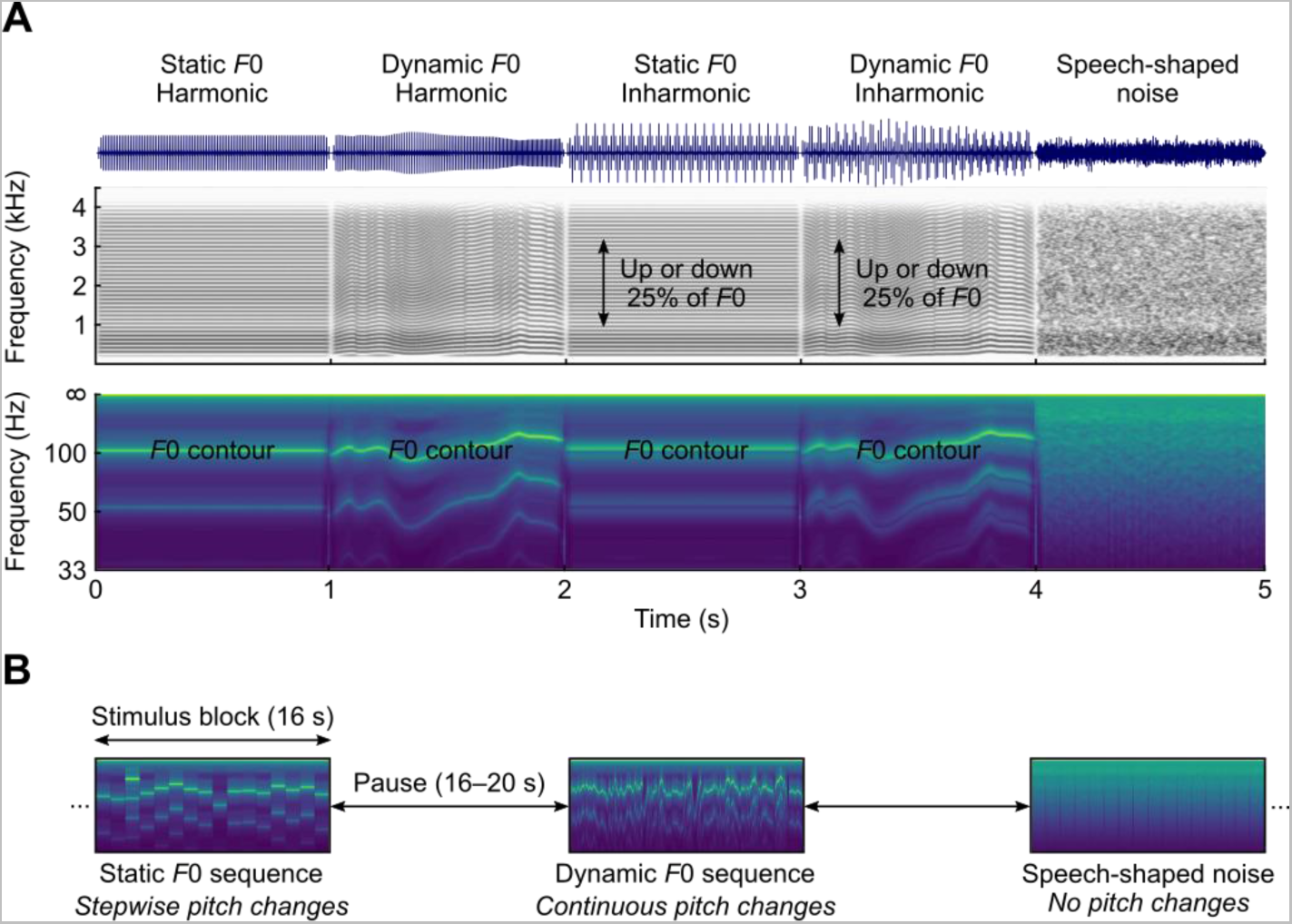
Example stimuli and experimental design. A) Waveforms, narrow-band spectrograms, and summary autocorrelation function (SACF) spectrograms showing the pitch contours for examples of the five stimulus types. B) The individual 1-s stimuli were presented as continuous blocks. Sequences consisting of static F0 stimuli were characterised by stepwise pitch changes between stimuli, those consisting of dynamic F0 stimuli exhibited continuous pitch changes. For speech-shaped noise sequences, there were no pitch changes at all.

### 2.3. Experimental design and procedure

The present experiment was originally designed as a simultaneous EEG and fNIRS study (Steinmetzger et al., 2020) and hence a block design was used to maximise the haemodynamic responses. Here, we re-analysed the EEG data, but omitted the fNIRS data because their limited depth resolution precludes fine-grained analyses of the activity emanating from deeper structures such as primary auditory cortex.

The individual 1-s stimuli in each condition were randomly concatenated into blocks of 16 stimuli and followed by pauses with random durations ranging from 16–20 s. Each participant was presented with 12 blocks of each of the 5 stimulus conditions, adding up to a total duration of about 34 mins. The order of the blocks was randomised without any constraints. As the EEG data were analysed relative to the onset of the individual 1-s stimuli in each block, this design resulted in 192 trials per condition. As shown in Fig. 1B, concatenating the harmonic or inharmonic stimuli with static *F*0s into blocks resulted in stepwise pitch changes at the onsets of the individual stimuli, while blocks consisting of harmonic or inharmonic stimuli with dynamic *F*0s were characterised by continuously changing, speech-like pitch contours. Blocks containing concatenated segments of speech-shaped noise were included as a control condition that had a similar spectral envelope but no pitch changes.

The experiment took place in a sound-attenuating and electrically shielded room, with the participant sitting in a comfortable reclining chair during data acquisition. There was no behavioural task, but pauses were inserted about every 10 mins to ensure the vigilance of the subjects. The stimuli were presented with 24-bit resolution at a sampling rate of 48 kHz using an RME ADI-8 DS sound card (Haimhausen, Germany) and Etymotic Research ER2 earphones (Elk Grove Village, IL, USA) connected to a Tucker-Davis Technologies HB7 headphone buffer (Alachua, FL, USA). The presentation level was set to 70 dB SPL using an artificial ear (Brüel & Kjær, type 4157, Nærum, Denmark) and a corresponding measurement amplifier (Brüel & Kjær, type 2610, Nærum, Denmark).

### 2.4. EEG recording and analysis

Continuous EEG signals were recorded using a BrainVision actiCHamp system (Brain Products, Gilching, Germany) with 60 electrodes arranged according to the extended international 10-20 system. Four additional electrodes were placed around the eyes to record vertical and horizontal eye movements. The EEG data were recorded with an initial sampling rate of 500 Hz, an online anti-aliasing low-pass filter with a cut-off frequency of 140 Hz and were referenced to the right mastoid. The electrode positions of each subject were digitized with a Polhemus 3SPACE ISOTRAK II system before the experiment.

The data were pre-processed offline in the same way as in Steinmetzger et al. (2020) using FieldTrip (version 20180924; Oostenveld et al., 2011) and custom MATLAB code. The continuous waveforms were first segmented into epochs ranging from −0.3–1.1 s around stimulus onset. Next, the epochs were re-referenced to the mean of both mastoids and detrended as well as demeaned by removing a 1^st^-order polynomial. The epochs were then low-pass filtered (cut-off 15 Hz, 4^th^-order Butterworth, applied forwards and backwards), baseline corrected by subtracting the mean amplitude from −0.1–0 s before stimulus onset, and subsequently down-sampled to 250 Hz. After visually identifying and excluding bad channels (total = 4, max. 2 per subject), the data were decomposed into 20 principal components to detect and eliminate eye artefacts. After the 4 eye electrodes were removed from the data, epochs in which the amplitudes between −0.2–1 s around stimulus onset exceeded ±60 µV or the *z*-transformed amplitudes differed by more than 15 standard deviations from the mean of all channels were excluded from further processing. On average, 86% of the trials (830/960 per subject, min. 65% per subject) passed the rejection procedure. Lastly, bad channels were interpolated using the weighted average of the neighbouring channels, the data were re-referenced to the average of all 60 channels, and again baseline corrected from −0.1–0 s before stimulus onset.

Distributed source reconstructions of the resulting event-related potentials (ERPs) were computed using the MNE-dSPM approach implemented in Brainstorm (version 10-Jun-2022; Dale et al., 2000; Tadel et al., 2011). The electrode positions of each subject were co-registered to the ICBM152 MRI template by first aligning three external fiducial points (LPA, RPA, and Nz) and subsequently projecting the electrodes to the scalp of the template MRI. A Boundary Element Method (BEM) volume conduction model based on the ICBM152 template and the corresponding cortical surface (down-sampled to 15,000 vertices) were used as head and source models. The BEM head model was computed using OpenMEEG (version 2.4.1; Gramfort et al., 2010) and comprised three layers (scalp, outer skull, and inner skull) with 1082, 642, and 642 vertices, respectively. Linear MNE-dSPM solutions with dipole orientations constrained to be normal to the cortex were estimated for each subject and condition after pre-whitening the forward model with the averaged noise covariance matrix calculated from the individual trials in a time window from −0.2–0 s before stimulus onset. The default parameter settings for the depth weighting (order = 0.5, max. amount = 10), noise covariance regularisation (regularise noise covariance = 0.1), and regularisation parameter (SNR = 3) were used throughout.

## 3. Results

### 3.1. Sensor-level ERPs

In a first step, the scalp ERPs evoked by stepwise and continuous pitch changes were compared after pooling together the harmonic and inharmonic versions of both conditions. As shown in Fig. 2A, stepwise pitch changes caused by transitions between stimuli with static *F*0s elicited markedly larger P2 amplitudes, as confirmed by a cluster-based permutation test (~160–352 ms, *t*_(cluster)_ = 4540.86, *p* < 0.001***, *d* = 1.52; Maris et al., 2007). This test was based on sample-wise dependent-samples *t*-tests with a cluster-forming threshold of *p* < 0.05 (two-sided), a minimum of 3 neighbouring electrodes per cluster, and 10,000 randomisations to determine the cluster *p*-values. The returned cluster had a fronto-central scalp distribution and included 24 electrodes at its midpoint (cf. scalp map insert in Fig. 2A).

**Figure 2.**
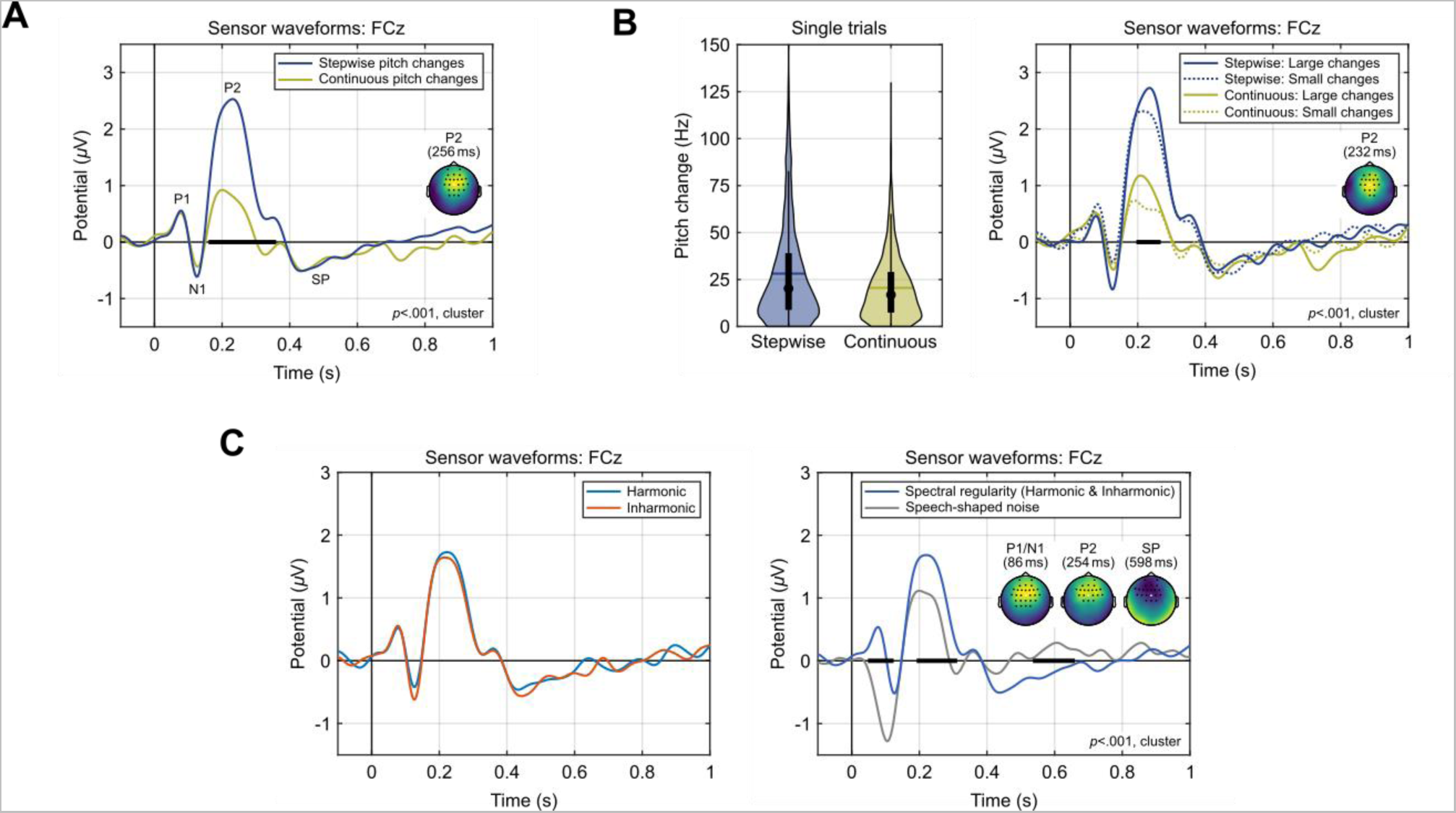
Sensor-level ERPs. Effects of pitch change type (A) and magnitude (B), as well as harmonicity and spectral regularity (C) on the P2 amplitude. ERPs traces are shown for electrode FCz, highlighted in the scalp maps. The thick horizontal black bars indicate significant time windows. In the scalp maps, the voltage of the second condition was subtracted from the first one, as indicated in the legends. The maps exhibit the voltage difference and the electrodes that were part of the respective cluster at its temporal midpoint. Violin plots in (B) show the distributions of pitch change magnitudes at the transitions between individual trials, along with ERPs after dividing the trials into subgroups with pitch change magnitudes above or below the overall median. SP = sustained potential.

To estimate the effect of the pitch change magnitude on the P2 amplitudes, the stimulus sequences for each individual participant were then re-constructed and the pitch steps between successive stimuli were calculated using the SACF software described above. As the average magnitude was found to be somewhat larger for stepwise pitch changes (means = 28.1/20.6 Hz; medians = 16.8/20.2 Hz; see Fig. 2B for the distributions), the single trials were divided into subgroups above and below the average median across both conditions (18.5 Hz). Additionally, trials in the stepwise condition were excluded from analysis if the magnitude exceeded the maximum of the continuous condition (129.6 Hz) to align the distributions. As illustrated in Fig. 2B, trials with a magnitude above the median elicited larger P2 amplitudes for both types of pitch change (~196– 268 ms, *t*_(cluster)_ = 1055.1, *p* < 0.001***, *d* = 1.61; data pooled across stepwise and continuous conditions for testing). However, the duration as well as the P2 amplitude difference of this effect were much smaller compared to the effect of pitch change type (cf. Fig. 2A).

Next, the ERP data were analysed regarding potential effects of harmonicity by comparing all harmonic and inharmonic stimuli (i.e., the stepwise and continuous pitch change conditions were pooled together). Fig. 2C shows that no such effects were evident throughout the entire duration of the stimuli and the largest cluster returned had a *p*-value of 0.537.

In contrast, when comparing all stimuli with a regular spectral structure (i.e., all harmonic and inharmonic conditions pooled together) to a control condition comprising speech-shaped noise, cluster-based testing indicated three separate highly significant clusters, as shown on the right side of Fig. 2C. These clusters were due to the absence of a P1 and the larger N1 elicited by speech-shaped noise (~48–124 ms, *t*_(cluster)_ = 3260.16, *p* < 0.001***, *d* = 2.25), and the increased P2 (~192–316 ms, *t*_(cluster)_ = 2254.94, *p* < 0.001***, *d* = 1.43) and sustained potential amplitudes (SP; ~536–660 ms, *t*_(cluster)_ = −2094.85, *p* < 0.001***, *d* = 1.33) evoked by stimuli with spectral regularity. All three clusters had a fronto-central scalp distribution and comprised at least 20 electrodes during the midpoint of the respective cluster time windows.

### 3.2. Source waveforms

The source waveforms were extracted from a set of pre-defined anatomical regions of interest (ROIs). As shown previously (Steinmetzger et al., 2020), the current stimuli mainly evoke activity in the supratemporal plane and the superior temporal sulcus (STS), as is typical for speech and speech-like stimuli (e.g., Belin et al., 2000). Hence, the set of ROIs chosen (Fig. 3A, top panel) comprised regions along the supratemporal plane [planum temporale (PT), Heschl’s sulcus (HS), Heschl’s gyrus (HG), and planum polare (PP)] as well as STS, bilaterally, as specified in the Destrieux atlas (Destrieux et al., 2010) implemented in Brainstorm. The underlying distributed source reconstructions, however, were computed across the entire cortical surface, without applying any ROI-based spatial restrictions.

**Figure 3.**
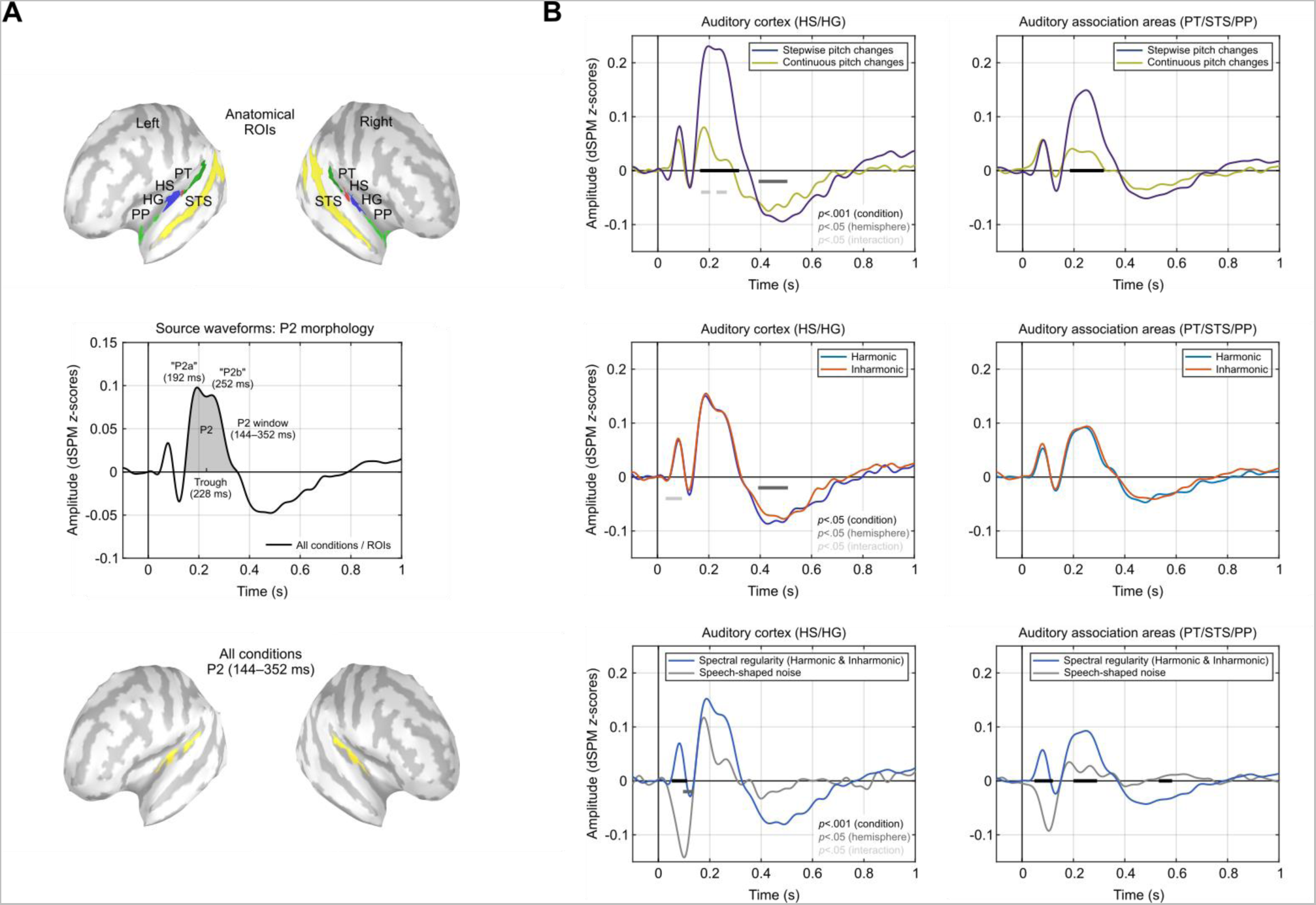
Source waveforms. A) Auditory anatomical regions of interest (ROIs) from which the source waveforms were extracted (top). Abbreviations: PT = planum temporale, HS = Heschl’s sulcus, HG = Heschl’s gyrus, PP = planum polare, STS = superior temporal sulcus. Two-peaked morphology of the P2 component after averaging across all stimulus conditions and ROIs (middle), and corresponding source localisation across the entire P2 window (bottom). B) Source waveforms for the contrasts of type of pitch change (top), harmonicity (middle), and spectral regularity (bottom). The waveforms are shown after averaging across ROIs belonging to auditory cortex and auditory associations areas and were averaged across hemispheres. The thick horizontal bars indicate significant time windows. Effects shorter than 25 ms were omitted throughout.

When averaged across all stimulus conditions and ROIs, the resulting source waveform was dominated by a large P2 (Fig. 3A, middle panel), same as the sensor waveforms. However, unlike the sensor-level ERPs, this source waveform exhibited a double-peaked morphology for the P2, with two separate peaks spaced approximately 60 ms apart (192/252 ms), hereafter referred to as “P2a” and “P2b”. We opted for a typical nomenclature based on morphology, where components are named based on their temporal sequence, unlike Benner et al. (2023) who referred to the anterior and posterior P2 sources as “P2” and “P2a”, respectively,

In a first step to identify the locations of the cortical generators of the P2, a source localisation across all stimulus conditions and the entire P2 window (144–352 ms) was then computed (Fig 3A, bottom panel). The areas showing the largest activity were located on the supratemporal plane in both hemispheres, consistent with the pre-defined set of ROIs. Here and in the remainder of the paper, the source maps were plotted such that activity beyond auditory areas was masked by adjusting the amplitude threshold and the minimum number of connected vertices accordingly.

Next, the source waveforms were averaged over ROIs belonging to auditory cortex (HG & HS) and auditory association areas surrounding auditory cortex (PT, STS & PP), and analysed using the same three condition contrasts as before (Fig. 3B). These contrasts were statistically evaluated via dependent-samples *t*-tests (two-sided) for each time point from 0–1000 ms, with *p*-values determined by permutation testing (10,000 randomisations). The reported *t*-values represent the average over the respective significant time window. To test for main effects of condition, the source waveforms were averaged across hemispheres, while main effects of hemisphere were evaluated by averaging across conditions. Interactions of condition and hemisphere were tested by comparing condition differences across hemispheres. Effects with a duration of less than 25 ms were considered as false positives and omitted throughout. In Fig. 3B, the source waveforms are depicted after averaging over ROIs and hemispheres, but the waveforms for each individual ROI and hemisphere are provided in Fig. 4.

**Figure 4.**
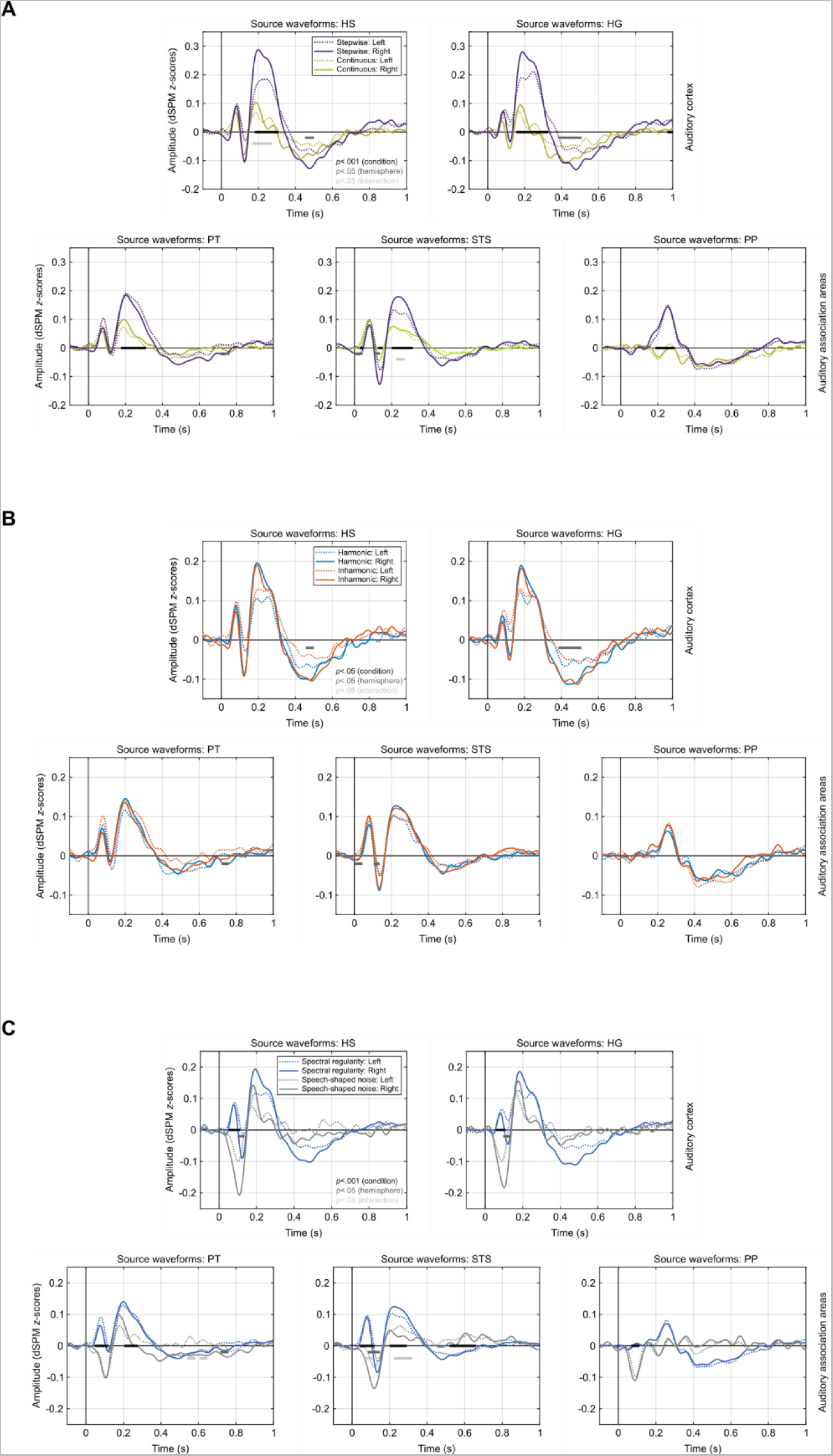
Source waveforms for the individual ROIs and hemispheres. The structure and details of the figure are the same as for the source waveforms shown in Fig. 3B.

The comparison of stepwise and continuous pitch changes (Fig. 3B, top row) showed that P2 amplitudes were significantly larger in both auditory cortex (164–320 ms, *t*_(19)_ = 4.94, *p* < 0.001***, *d* = 1.26) and association areas (184–316 ms, *t*_(19)_ = 4.93, *p* < 0.001***, *d* = 1.23) following stepwise changes. In auditory cortex, a main effect of hemisphere (392–504 ms, *t*_(19)_ = 2.61, *p* < 0.05*, *d* = 0.61) furthermore indicated that the sustained potential was overall larger in the right hemisphere. In addition, the particularly large P2 amplitudes elicited by stepwise pitch changes in right auditory cortex (Fig. 4A) resulted in a significant condition*hemisphere interaction (168–204 & 228–268 ms, *t*_(19)_ = 2.39, *p* < 0.05*, *d* = 0.61).

For the harmonicity contrast (Fig. 3B, middle row), there were again no significant condition differences (*p* < 0.05) for the P2 or any other component, neither in auditory cortex nor association areas. Even when considering all ROIs separately, no significant main effects of condition were evident (Fig. 4B). However, a main effect of hemisphere (392–504 ms, *t*_(19)_ = 2.54, *p* < 0.05*, *d* = 0.60) indicated a larger sustained potential in right auditory cortex, and a larger P1 for the inharmonic condition in left auditory cortex resulted in a significant interaction (36–92 ms, *t*_(19)_ = 2.49, *p* < 0.05*, *d* = 0.67).

The contrast of spectral regularity and speech-shaped noise (Fig. 3B, bottom row), on the other hand, revealed highly significant condition differences during the P1/N1 period in both auditory cortex (52–112 ms, *t*_(19)_ = 8.96, *p* < 0.001***, *d* = 2.25) and association areas (48–120 ms, *t*_(19)_ = 8.23, *p* < 0.001***, *d* = 2.16), as well as during the P2 (200–292 ms, *t*_(19)_ = 5.39, *p* < 0.001***, *d* = 1.35) and SP windows (532–584 ms, *t*_(19)_ = 3.71, *p* < 0.001***, *d* = 0.86) in association areas. In line with the sensor-level results, P1, P2, and SP were thus larger in amplitude for the stimuli with spectral regularity, while the N1 was enhanced for speech-shaped noise. In addition, the source-level results showed that the P2 and SP effects emerged from auditory association cortex, particularly STS (Fig. 4C). Furthermore, a main effect of hemisphere was observed for the N1 (96–136 ms, *t*_(19)_ = 2.59, *p* < 0.05*, *d* = 0.61), indicating larger amplitudes in the right auditory cortex in both conditions.

### 3.3. Cortical generators of the P2 subcomponents

In the second part of the source-level analyses, distributed source reconstructions were used to determine the cortical generators of the P2a and P2b subcomponents observed in the source waveforms. Since the largest P2 amplitudes were evoked by stepwise pitch changes, we first evaluated the source maps for these stimuli. As shown in Fig. 5A, the time-averaged source activity for the P2a (144–228 ms) was greatest in right auditory cortex, whereas the generators of the P2b (228–352 ms) were more broadly distributed along the supratemporal planes in both hemispheres. The time windows of the P2 subcomponents were derived by dividing the entire P2 window (144–352 ms) of the grand-average source waveform across all conditions and ROIs into segments before and after the trough at 228 ms that separates the two P2 peaks (cf. Fig. 3A).

**Figure 5.**
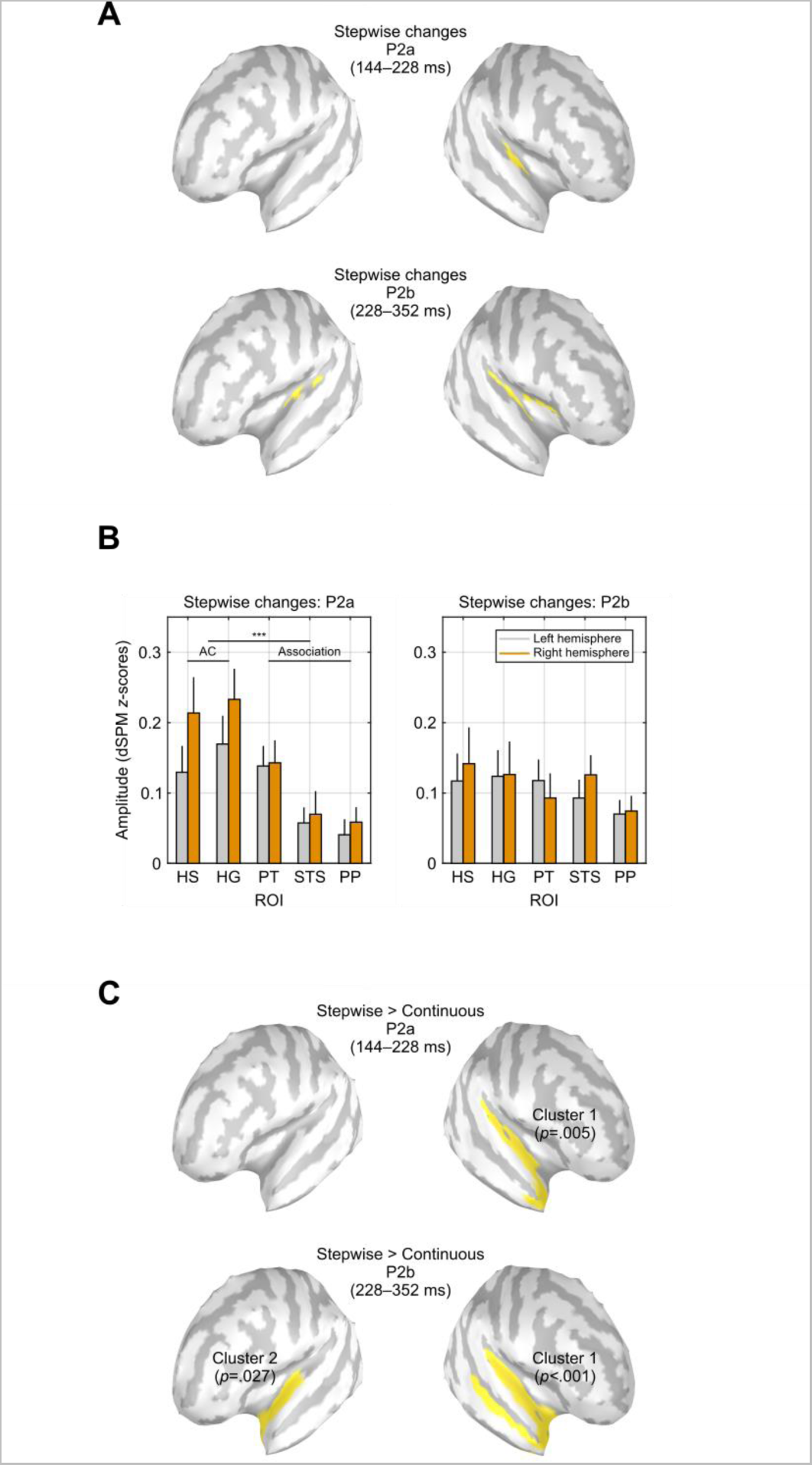
Source analysis of the P2 subcomponents. A) Cortical generators of the P2a and P2b subcomponents elicited by stepwise pitch changes. B) Source amplitudes of the P2a and P2b elicited by stepwise pitch changes, separately for each auditory ROI and hemisphere. C) Statistical comparison of the P2a and P2b source reconstructions for stepwise and continuous pitch changes. Yellow regions indicate auditory areas where stepwise changes evoked significantly greater activity.

In Fig. 5B, the averaged source amplitudes in response to stepwise pitch changes are shown for each ROI and hemisphere. As confirmed by a repeated-measures ANOVA, for which the activity was again averaged across ROIs belonging to either auditory cortex or auditory association cortex, activity for the P2a time window was significantly greater in auditory cortex compared to association areas (*F*_(1,19)_ = 29.2, *p* < 0.001***, η^2^ = 0.61). Driven by the stronger activity in right auditory cortex, there was also a significant interaction of region and hemisphere (*F*_(1,19)_ = 6.59, *p* = 0.019*, η^2^ = 0.26). However, a post-hoc *t*-test showed that activity in the right and left auditory cortex did not differ significantly despite a strong trend (*t*_(19)_ = 1.95, *p* = 0.067, *d* = 0.44). On the contrary, neither main effects nor an interaction was observed for the P2b window (*p ≥* 0.119), indicating a comparable level of activity across auditory areas.

Finally, the distributed source reconstructions of the P2 subcomponents evoked by stepwise and continuous pitch changes were statistically compared (Fig. 5C). To identify auditory regions where stepwise changes elicited greater activity than continuous changes, cluster-based permutation tests were computed, for which the source amplitudes of each vertex on the cortical surface were averaged over the respective P2 time window. These tests were based on dependent-samples *t*-tests for each vertex with a cluster-forming threshold of *p* < 0.05 (one-sided), a minimum of 3 neighbouring vertices per cluster, and 10,000 randomisations to determine the cluster *p*-values. Only clusters overlapping with the auditory ROIs are reported (cf. Fig. 3A).

For the P2a window, a single significant cluster (*t*_(cluster)_ = 890.98, size = 192 vertices, *p* = 0.005**, *d* = 1.77) indicated that activity in response to stepwise pitch changes was stronger across the length of the right supratemporal plane. For the P2b window, two separate clusters indicated greater activity following stepwise changes along the right supratemporal plane as well as STS (*t*_(cluster)_ = 1081.39, size = 319 vertices, *p* < 0.001***, *d* = 1.07) and the anterior portion of the left supratemporal plane (*t*_(cluster)_ = 457.94, size = 155 vertices, *p* = 0.027*, *d* = 0.90). Despite the greater spatial extent, the distribution of these clusters is in line with the locations of the generators of the P2 subcomponents observed in response to stepwise pitch changes shown in Fig. 5A. Whereas the first P2 subcomponent mainly originated from auditory cortex, particularly in the right hemisphere, the second subcomponent showed a broader distribution including auditory and association areas in both hemispheres.

## 4. Discussion

### 4.1. Functional significance of the P2 and its subcomponents

In summary, the present results have shown that the P2 amplitude reflects the type and, to a lesser extent, the magnitude of voice pitch changes in sequences of speech sounds. The P2 was the only auditory ERP component affected by these two factors, suggesting that it is the primary index reflecting the cortical processing of spectrally complex sounds such as speech. The two types of voice pitch changes employed comprised stepwise and continuous changes. In sequences with stepwise pitch changes, the individual sounds had a static pitch resembling monotonised speech and pitch changes were consequently restricted to the transitions between sounds. In contrast, sequences with continuous pitch changes were formed of sounds with dynamically varying pitch contours extracted from natural speech. Hence, there were pitch jumps between the individual stimuli as well as continuous pitch changes throughout the sequences. In both sequence types, other acoustic factors such as duration, level, and spectral envelope were kept constant. The substantially larger P2 evoked by stepwise changes shows that the context in which voice pitch changes occur is the crucial factor determining the P2 amplitude. The driving factor behind this effect appears to be the greater saliency of the stepwise pitch changes. The ongoing modulation of the pitch contours in sequences with continuous pitch changes likely resulted in a greater degree of neural adaptation compared to sequences with stepwise pitch changes.

As is evident from the source waveforms (cf. Fig. 3B), both types of pitch changes as well as speech-shaped noise evoked a discernible P2a in auditory cortex. Although the amplitude of this peak was much larger for stepwise pitch changes compared to the other two conditions, the first P2 subcomponent is thus elicited irrespective of the acoustic properties of the materials. The P2a might therefore represent an obligatory initial processing step indicating that some form of change to the pitch or spectral structure of the stimuli has occurred. The second P2 subcomponent, in turn, was practically absent for all conditions except stepwise pitch changes, both in auditory cortex and association areas. Hence, it could be that a sufficiently large first P2 subcomponent triggers additional processing on the next level of the cortical hierarchy, as reflected in the P2b. The assumption that P2a and P2b are generated at different stages of the cortical processing hierarchy is supported both by their temporal sequence and the respective source localisations, which showed that the P2b emanated from areas located further away from primary AC than the P2a. For example, the cortical generators of the P2b included STS, a region exhibiting voice-selective activity (Belin et al., 2000), suggesting that the stimuli were classified as voice-like at this point. Thus, the P2b may indicate more detailed processing to identify the specific type of sound and in what way it changed.

### 4.2. Cortical generators of the P2 and its subcomponents

The distributed ERP source reconstruction computed across all stimulus conditions and the entire P2 window revealed bilateral foci of activity in PT and around HG (cf. Fig 3A). This finding agrees with the depth-electrode recordings (Godey et al., 2001) as well as fMRI-based MEG source localisations (Benner et al., 2023) suggesting that there might be separate cortical sources of the P2 in PT and anterior to primary auditory cortex. Dipole-based source localisations of MEG data, in contrast, have consistently reported only a single P2 generator in the lateral part of HG (Hari et al., 1987; Pantev et al., 1996; Ross et al., 2009; Tiitinen et al., 1999). The PT has been argued to be the cortical site in which complex spectro-temporal patterns in auditory scenes are segregated and compared with learned representations (Griffiths et al., 2002). Hence, its involvement in the processing of the spectro-temporal modulations constituting voice pitch changes, as reflected by the P2, appears plausible from a functional point of view. Furthermore, dipole sources in PT have recently been reported for the P2 evoked by voice pitch changes in an EEG study with unilateral CI users with preserved contralateral hearing, regardless of which ear received auditory input (Steinmetzger et al., 2022b).

Based on the particularly large P2 evoked by stepwise voice pitch changes, which was assumed to allow for robust estimations of the underlying cortical sources due to its favourable signal-to-noise ratio, we then examined if the two separate P2 peaks evident in the source waveforms are generated in different cortical areas. For the first subcomponent, termed P2a, activity in the auditory cortex was significantly stronger than in the surrounding auditory association cortex and there was a pronounced trend for more activity in the right hemisphere, in line with a previously reported right lateralisation of the P2 in response to speech (Coffey et al., 2017) and voice pitch changes (Steinmetzger et al., 2022a). Furthermore, pitch changes in speech and music have been shown to be processed in the anterior portion of the right auditory cortex in several neuroimaging and lesion studies (e.g., Johnsrude et al., 2000; Patterson et al., 2002; Zatorre et al., 2001). For the subsequent P2b, in contrast, a comparable degree of activity was evident across all auditory ROIs and activity levels were similar across hemispheres. The wide network of cortical regions involved in generating the first and particularly the second P2 subcomponent suggests that distributed source reconstructions might be better suited to identify the cortical generators of the P2 than classic dipole solutions.

Regarding the choice of auditory ROIs and their allocation to auditory cortex and auditory association cortex, respectively, we opted for a simple scheme that takes the limited spatial resolution of EEG source reconstructions into account. Thus, a rather coarse macro-anatomical atlas that only distinguishes between gyri and sulci was used (Destrieux et al., 2010). However, as there is no strict correspondence between macro-anatomy and cytoarchitecture, a consensus concerning the organisation of the human auditory cortex is still lacking (Moerel et al., 2014; Zachlod et al., 2020). For example, according to Moerel et al. (2014), the majority of PT is part of the parabelt and may thus be considered part of auditory cortex, whereas Griffiths et al. (2002) noted that it is “generally agreed to represent auditory association cortex”. For simplicity, we hence distinguished between a smaller core region taken to comprise primary and secondary auditory cortex (“auditory cortex”; HG & HS) and a surrounding larger region consisting of areas associated with higher-order auditory processing (“auditory association cortex”; PT, STS & PP).

### 4.3. Spectral regularity rather than harmonicity drives the P2

The current results revealed no differences in P2 amplitude between harmonic stimuli and their inharmonic equivalents. This finding applies to all auditory cortical regions examined as well as all other ERP components besides the P2. When pooling the harmonic and inharmonic conditions together, however, the P2 was markedly larger compared to a control condition of speech-shaped noise that had no spectral regularity, i.e., no discrete spectral components. The latter finding is in line with fMRI results showing enhanced responses to harmonic sounds compared to spectrally-matched noise across human auditory cortex (Norman-Haignere et al., 2013; Norman-Haignere et al., 2019).

The stimuli were rendered inharmonic by shifting all spectral components in frequency, a technique that maintains the presence and regular spacing of spectral components and leaves the envelope modulations unaffected. These properties, which have recently been verified by detailed acoustic analyses (Steinmetzger et al., 2023), make this stimulus type ideally-suited for investigating potential effects of stimulus harmonicity. The same psychoacoustic study also showed that the intelligibility of a target speech signal masked by harmonic or frequency-shifted inharmonic tone complexes did not differ, in agreement with the current findings. In the neurosciences, however, shifted inharmonic stimuli have previously only been used in animal studies. Invasive recordings from marmosets (Feng et al., 2017; Wang, 2018) and rabbits (Su et al., 2020) have provided evidence for the existence of so-called “harmonic template neurons” that show increased firing in response to harmonic sounds. Yet, at least in the core auditory cortex of marmosets (Feng et al., 2017), there appear to be relatively few such neurons. Assuming the same applies to the human auditory cortex, the current results suggest that it may not be possible to detect the responses of these neurons in non-invasive recordings due to their limited number.

The absence of an effect of harmonicity in the current study furthermore implies that the pitch strength of the stimulus materials *per se* does not affect auditory cortex activity. As can be seen in Fig. 1A, and demonstrated in more detail in Steinmetzger et al. (2023), the pitch strength, or periodicity, of the inharmonic stimuli is markedly lower than that of their harmonic equivalents. This makes for a different reading of experiments that have investigated pitch-related responses in auditory cortex. Several of the studies that reported enhanced activity in the ‘pitch centre’ located at the anterolateral border of HG in response to sounds giving rise to a pitch percept contrasted pulse trains with regular and irregular spacing (Bendor et al., 2005; Bendor et al., 2010; Gutschalk et al., 2011; Gutschalk et al., 2002; Gutschalk et al., 2004). Yet, while irregular pulse trains do not evoke a clear pitch, this manipulation also results in stimuli without discrete spectral components. The same is true for studies using iterated rippled noise (IRN; Griffiths et al., 1998; Ritter et al., 2005). When the number of iterations in the construction of IRN materials is reduced, both the pitch and spectral peaks dissipate. Lastly, the pitch strength of the materials has also been altered by either reducing the number of resolved harmonics (Norman-Haignere et al., 2013) or by using complex tones with only resolved or unresolved harmonics, respectively (Penagos et al., 2004). Crucially, none of the stimuli used in the above studies allowed for a manipulation of the pitch strength that is independent of the presence and number of spectral components, unlike the shifted inharmonic materials used in the present experiment. In agreement with this, Feng et al. (2017) also did not report increased responses in the marmoset pitch centre for harmonic compared to shifted inharmonic stimuli. It is thus conceivable that the activity in the cortical pitch centre merely reflects the number of discrete, regularly-spaced spectral components in the stimulus materials rather than their pitch.

## Data availability

The stimuli and EEG data are available at https://osf.io/tnfdg and the code used to process the data is available at https://osf.io/bnzmy.

## Abbreviations

CI: Cochlear implant
EEG: Electroencephalography
ERPs: Event-related potentials
*F*0: Fundamental frequency
fNIRS: Functional near-infrared spectroscopy
HG: Heschl’s gyrus
HS: Heschl’s sulcus
MEG: Magnetoencephalography
PP: Planum polare
PT: Planum temporale
ROI: Region of interest
SACFs: Summary autocorrelation functions
SP: Sustained potential
STS: Superior temporal sulcus.

